# Dynamic Culture Improves the Predictive Power of Bronchial and Alveolar Airway Models of SARS-CoV-2 Infection

**DOI:** 10.1101/2025.07.21.665885

**Authors:** Claire H Caygill, Rose Lopeman, Kirstie A Lewis, Emily Richardson, Aitor Casas Sanchez, Nick Heavey, Adam Winrow, Louise Howard, Chris Williams, Domonic Wooding, Thomas Edwards, Eric Lucas, Tomasz Kostrzewski, Andrew Owen, Shaun H Pennington, Giancarlo A Biagini

**Affiliations:** Centre for Drugs and Diagnostics, Department of Tropical Disease Biology, Liverpool School of Tropical Medicine, Pembroke Place, Liverpool, L3 5QA, United Kingdom; CN Bio, Cambridge Science Park Milton Road, Milton, Cambridge, CB4 0WN, United Kingdom; Department of Pathology, University of Cambridge, Cambridge, CB2 1QP; Department of Pharmacology and Therapeutics, Centre of Excellence in Long-acting Therapeutics (CELT), University of Liverpool, Liverpool, L69 3GE, United Kingdom

## Abstract

Human *in vitro* lung models represent advanced tools for studying respiratory infections, particularly those caused by emerging respiratory pathogens. Despite scientific advances, vaccine and therapeutics pre-clinical development has yet to fully adopt human-relevant testing platforms due in part to a lack of validation. In this study, we characterised how static and dynamic flow culture conditions influence microphysiological systems (MPS) generated using primary bronchial and alveolar epithelial cells. We assessed epithelial structure, functional differentiation, and infection dynamics. This study represents the first direct comparison of how dynamic flow and endothelial co-culture influence viral tropism, replication kinetics, and host responses across anatomically distinct regions of the respiratory tract *in vitro*.

Dynamic flow promoted formation of more physiologically relevant tissue architecture, pseudostratified bronchial epithelium and alveolar sac-like structures, with enhanced epithelial differentiation and retention of region-specific cell phenotypes at the transcriptomic level. Both static and dynamic flow models demonstrated responsiveness to inflammatory stimuli (poly(I:C), LPS), producing distinct, tissue-specific cytokine profiles and supporting infection with multiple SARS-CoV- 2 variants. Differences in infection efficiency, viral replication, and host gene expression were observed between variants, with dynamic flow models offering enhanced sensitivity and resolution. In alveolar tissues, dynamic flow increased infection efficiency and reduced variability, enabling more robust and consistent transcriptional responses. This facilitated the identification of interferon signalling pathways as key targets of the host response. Among the variants tested, Delta induced the most extensive tissue damage and strongest transcriptional response, whereas Omicron BA.5 exhibited greater infectivity in alveolar models compared to earlier variants.

Our findings demonstrate that dynamic flow MPS more closely replicate human lung tissue architecture and cellular diversity, while also enhancing the predictive power and clinical relevance of airway models for *ex vivo* studies of SARS-CoV-2 infection. These improvements strengthen the reliability of data generated for the study of host–pathogen interaction studies and support the use of dynamic systems for evaluating novel anti-infectives, immunomodulators, and functional characterisation of immune sera generated by next-generation vaccines. Collectively, our results highlight the value of integrating dynamic in vitro models into preclinical pipelines for emerging respiratory pathogens.

## Introduction

Microphysiological systems (MPS) represent a rapidly advancing field, incorporating technologies such as organ-on-a-chip (OOC) and multi-cellular 3D organoids, with the aim of more accurately replicating the complex physiological environment observed in vivo. A range of lung-on-a-chip (LOC) platforms - such as those developed by CN Bio, Emulate, AlveoliX and PREDICT96 - have demonstrated promising physiological relevance. However, a recent meta-analysis comparing OOC and static models reported only modest improvements in biomarker expression within perfused tissue. In addition, the meta-analysis highlighted the paucity of data comparing dynamic models studies that have benchmarked against static model against their static counterparts. This lack of comparative data is a key barrier to the broader adoption of MPS as alternatives to animal models in the preclinical development of vaccines and therapeutics [5]. The issue is particularly evident in infectious disease research, where multiple studies have demonstrated that MPS can support replication of respiratory pathogens [6–9], but, to our knowledge, none have formally validated these models against conventional 2D/static systems to determine whether their added complexity translates into improved functional performance.

Further to this, there remains a need for robust and reproducible models that accurately represent distinct regions of the respiratory tract, as clinically relevant differences in cell type composition and tissue morphology exist between these sites [9]. Understanding infection dynamics and host–pathogen interactions across different airway regions is critical to elucidate how respiratory infections progress from the upper airway into the lower alveolar region, where patients are at greater risk of complications such as acute respiratory distress syndrome (ARDS). Modelling the full respiratory tract also provides a platform to monitor viral evolution and may improve clinical risk assessment for emerging pathogens.

To address this gap, we developed and characterised both bronchial and alveolar airway models using static Transwell® cultures and CN Bio’s lung-on-a-chip PhysioMimix® system (hereinafter referred to as ‘dynamic flow MPS’). Each model incorporated primary human airway epithelial cells and pulmonary endothelial cells to better replicate native lung architecture. We evaluated tissue morphology, cell type differentiation, and physiological function - including barrier integrity and inflammatory responses - following infection with early and emerging SARS-CoV-2 variants: Pre-alpha, Delta (B.1.617.2), and Omicron (BA.5). Infection efficiency, viral replication kinetics, and host responses were assessed across both static and dynamic systems. SARS-CoV-2, a beta coronavirus and the causative agent of COVID-19, primarily targets the respiratory tract, leading to a spectrum of disease from asymptomatic infection to severe ARDS and death. Improved understanding of early-stage infection and regional airway responses is essential for guiding the development of vaccines and targeted therapeutics. As such, SARS-CoV-2 provided an ideal model pathogen to evaluate the performance of static versus dynamic flow airway infection platforms.

Here, we demonstrate that dynamic flow culture improves tissue architecture and cellular differentiation relative to static systems. We also show variant- and airway-specific differences in SARS-CoV-2 infection dynamics and host responses, highlighting the utility of lung MPS platforms for infectious disease research and preclinical evaluation of therapeutic interventions.

## Methods

### Generation of static and dynamic flow MPS coculture bronchial and alveolar airway models

Airway models were comprised of human pulmonary microvascular endothelial cells (HPMEC) (PromoCell, C-12281, lot.489Z024.1) and small airway epithelial cells (SAEC) (Lonza, CC-2547, lot.21TL214918) or normal human bronchial epithelial cells (NHBE) (Lonza, CC-2540, lot.21TL035497) for the alveolar and bronchial airway model, respectively. HPMECs, SAECs and NHBEs were maintained at 37°C, 5% CO^2^ and 95% humidity in cell-type specific media (EGM-2 basal medium bullet kit [Lonza, CC-3162]; small airway epithelial cell media [PromoCell, C-211170]; PneumaCult-Ex Plus medium [STEMCELL Technologies, 05040], respectively) in T75 flasks until they reached 70% confluency. Cell media was changed every 48-72 hrs. Once at 70% confluency, cells were washed with PBS and incubated with TrypLE (Gibco; 14170088) at 37°C to facilitate detachment. After 5 minutes, cell media was added, cells recovered and centrifuged at 300 × g for 5 minutes. The supernatant was then removed, and the cells resuspended in cell media. Cells were counted and seeded onto new T75 flasks at 1x10^6^ cells/mL and grown until required for airway model generation.

All airway models were produced in 24-well, 6mm, 0.4µM pore, PET Transwell® membranes (Corning; 3470) coated with 10 µg/mL type I rat tail collagen (Corning; 354236) in PBS at 37°C for 1 hour. After which, Transwells® were washed three times with PBS and stored at 4°C for up to two weeks. Prior to use, Transwells® were inverted and warmed to 37°C and HPMEC cells seeded at 5x10^4^ cells per well onto the basolateral surface in 75 µL seeding media (consult CN Bio, Cambridge UK for formulation). Transwells® were then returned to the incubator for 2 hrs to allow cell attachment. Transwells® were then transfered into either 24 well culture plates (static model) or MPS-T12 plates (CN Bio, Cambridge, UK) containing 600 µL seeding media in each well. SAEC or NHBEs were then seeded onto the apical surface at 1x10^5^ cells per well in seeding media. Static plates were returned to the incubator and MPS-T12 plates were placed in the PhysioMimix® docking station, without flow and incubated at 37°C.After 24 hours, MPS-T12 plates were subject to media flow at 0.5 µL/s.

48 hours after seeding, static and MPS-T12 plates were introduced to an air-liquid interface (ALI) by removal of media from the apical surface; media on the basolateral side was also removed and replaced with 600 µL differentiation media (consult CN Bio, Cambridge UK for formulation). Static plates were then again returned to the incubator and MPS-T12 plates placed back in the docking station with a flow rate of 0.5 µL/s to replicate human blood flow [10] for 14-16 days. The basolateral differentiation media was changed every 48-72 hours. At the same time, the apical surface was washed with 200 µL PBS to remove accumulated mucus debris.

### Assessment of Cellular Barrier Integrity

To ensure cell barrier integrity and development of tight junctions, transepithelial electrical resistance (TEER) measurements were performed every 48-72 hours whilst conducting apical surface washes using a Millicell® ERS Voltohmmeter (Merck, Darmstadt, Germany). TEER values were corrected by deducting the background TEER values measured from Transwell® inserts with no cells seeded and media/PBS only. TEER values were then multiplied by membrane surface area to calculate Ω cm^2^.

### LPS/poly(I:C) challenge

Twenty one days after seeding, media were changed and apical surface of the tissues incubated with PBS for 30 mins. Following removal of PBS, 50 µL of lipopolysaccharide (LPS; L4391, Merck) at 0.01 µg/mL or polyinosinic:polycytidylic acid (poly(I:C); 4287, BioTechne) at 10 µg/mL was added to the apical side of the Transwell® and media samples taken at 0, 2-, 6-, 24- and 48-hours following challenge. IP-10 release was analysed using Human CXCL-10/IP-10 DuoSet ELISA (DF1900, R&D Systems).

### Infection with pseudotyped SARS-CoV-2 virus

Fourteen days after seeding, tissues were media changed and washed on the apical side of the Transwell® with PBS for 30 mins at 37°C before being removed. SARS-CoV-2_S (D614G) pseudotyped lentivirus (EGFP) (P210906-1019gzb, VectorBuilder) was diluted in HBSS to obtain an MOI of 100 and 10% polybrene added at a final volume of 20 µL per well. The pseudotyped SARS-CoV-2 virus was added to the apical surface and incubated at 37°C for 48 hours, after which the inoculum was removed, and cultures were maintained under ALI conditions. Seven days post-infection, cells were fixed with 4% PFA for 30 minutes, followed by 4 × PBS washes and stored at 4°C for later use.

### SARS-CoV-2 Strains

SARS-CoV-2/Human/Liverpool/REMRQ0001/2020 was used for this work as an early reference strain and is referred to as ‘Pre-alpha’ for the remainder of this article. This isolate was collected from a nasopharyngeal swab from a patient in Liverpool. The mapped RNA sequence has previously been submitted to Genbank, accession number MW041156.

Delta (B.1.617.2) was gifted from Dr Thomas Edwards from Liverpool School of Tropical Medicine. Omicron (BA.5) was gifted from Professor James P Stewart from University of Liverpool.

All SARS-CoV-2 variants were propagated a maximum of 4 times in Vero E6 cells for downstream applications and variants were confirmed by nanopore sequencing (supplementary file 1).

### SARS-CoV-2 Infection and sample collection

Fourteen to sixteen days after maturation MPS-T12 plates were removed from the PhysioMimix® docking station and driver and transferred, along with static plates, to an incubator at 37°C, 5% CO^2^ and 95% humidity in the containment level 3 laboratory. SARS-CoV-2 variants were diluted in differentiation medium to obtain an MOI of 1 or 0.01 for a final volume of 70 µL per well. SARS-CoV- 2 variants or differentiation media only controls were added to the apical surface and incubated at 37°C for two hours. After two hours, the apical surface was washed 4 × with 200 µL PBS and the final wash retained as the day 0 sample. The basolateral side was washed 2 × with 600 µL PBS and 600 µL fresh differentiation media was then added. Cells were then returned to the incubator for 48-72 hours. On day 2, 4 and 7, differentiation media was replaced, and the apical surface washed twice with 100 µL PBS. PBS washes were pooled and retained for downstream processing. Seven days post-infection, cells assigned for immunofluorescent staining were fixed with 4% PFA for 30 minutes, followed by 4 × PBS washes and stored at 4°C for later use. Remaining lung cell samples were RNA extracted for use with NanoString gene expression analysis.

### Quantification of SARS-CoV-2 RNA

RNA was extracted from apical wash supernatants using the Qiagen QIAcube HT® and RNeasy® 96 QIAcube HT Kit (Qiagen, Germantown, USA) as per the manufacturer’s instructions. SARS-CoV-2 viral RNA was then detected using Coronavirus COVID-19 genesig® Real-Time PCR assay (PrimerDesign™ Ltd, Eastleigh, UK), as per manufacturer’s instructions on a QuantStudio™ 5 Real- Time PCR System (ThermoFisher Scientific). SARS-CoV-2 RNA copies/µL were calculated from serial dilutions of known SARS-CoV-2 positive controls.

### Quantification of live SARS-CoV-2

SARS-CoV-2 variant stock and experimental supernatant concentrations were determined by plaque assay in Vero E6 cells. Pre-seeded Vero E6 cells were incubated with serially diluted viral supernatants in Eagle’s Minimum Essential Medium (EMEM [Gibco]) in triplicate. Semi-solid media (EMEM supplemented with 4% HI FBS and 0.1% agarose) was then added to each well in a 1:1 ratio and plates were incubated at 37°C with 5% CO^2^ for 72 hours. After which, paraformaldehyde (PFA) was added to each well to achieve a final concentration of 4% and the plate incubated for 30 minutes at room temperature. To visualise the plaques, the medium was removed, cells were stained with crystal violet (70% v/v H2O, 10% v/v ethanol, 20% v/v methanol and 0.25% crystal violet powder [Sigma Aldrich]) and washed three times with water. The number of plaques in each well were enumerated at the highest countable concentration and the average value was used to calculate the titers in viral plaque forming units (PFU)/mL.

### Immunofluorescence staining

Lung tissues were washed with PBS before fixation with 4% paraformaldehyde (Alfa Aesar; J61899) for 30 minutes. Following a further 3 PBS washes the cells were either processed for histology or fixed and stained for immunofluorescence analysis. To obtain slices of tissue, samples were processed by the histology core facility at the University of Cambridge Pathology Department. Briefly, following membrane excision from the Transwell and storage in 70% ethanol, the membranes were placed on lens paper, placed in an embedding cassette and submerged in 70% ethanol. The cassettes were then placed in increasing concentrations of ethanol for 10 mins (70%, 80%, 95%, 100%), followed by two changes of xylene as a clearing solvent. Following removal from the cassette, the membranes were infiltrated with paraffin for 30 mins and halved using a microtome blade. With the cut side down, the membranes were embedded in paraffin and the cassette placed on top to cool in ice. The block was then sectioned at 5 µm slices, collected on a positively charged glass slide and baked at 60 °C for 1 hr. Tissues were stained using haematoxylin and eosin (H&E) for general morphology, and Alcian Blue was used separately to visualise acid mucins in bronchial cultures.

For immunofluorescence staining, cells were permeabilised using 0.3% Triton-X-100 (Merck; 108643) in PBS for 5 mins before being washed 3 times in 0.1% Tween-20 (Merck; P9416) in PBS (PBST). Tissues were then incubated for 1 hr in blocking buffer (1% w/v BSA, 0.5% goat serum, 0.5% FBS in PBST (Merck; A0281, G9023, Gibco; 10500064)). The tissues were then incubated with primary antibodies for 1 or 2 hr at room temperature in blocking buffer. Bronchial tissues were stained for: anti-acetylated α-tubulin (1:2000, Merck; T7451), anti-MUC5AC (1:1000, Merck; HPA040615). Alveolar cultures were stained for: anti-RAGE/AGER (1:500, Invitrogen, MA5-29007), anti-SFTPB (1:1000, ThermoFisher, PA5-42000). SARS-CoV-2 infected and uninfected control tissues were stained for anti-SARS-CoV-2 nucleocapsid protein (1:2000, Abcam, ab280201).

Cultures were then washed three times with PBST and incubated with secondary antibodies at 1:200 for 1 hr in the absence of light at room temperature, including: donkey anti-mouse Alexa Fluor 488 (abcam, ab150109), goat anti-rabbit Alexa Fluor 488 ([1:1000 dilution with anti-SARS-CoV-2] abcam, ab150077), goat anti-rabbit Alexa Fluor 594 (abcam, ab150080), goat anti-rabbit Alexa Fluor 647 (abcam, ab150079), goat anti-mouse Alexa Fluor 647 (abcam, ab150115) and co-stained with Phalloidin-FITC or -TRITC (1:200, abcam, ab235137 or Merck, P1951 respectively) and Hoechst 33342 (1:5,000, Invitrogen, H3570) or DAPI (1:1000, Invitrogen, D1306). Cultures were washed three times with PBST, membranes were excised from the Transwells® and mounted onto glass coverslides for imaging. Z-slices were obtained using the Nikon A1-R point-scanning confocal microscope and analysed using ImageJ/Fiji. Infected tissues and uninfected control images were acquired using Zeiss LSM 880, Axio Observer, confocal microscope at 40x objective with camera (Zeiss, Germany) located in the CL3 facility and analysed with Zen Blue software.

### Cell Lysate RNA extraction

Seven days post-infection lung cells on the apical surface of Transwells® were lysed with 500 µL TRIzol™, scraped off the membrane surface and transferred to Phasemaker™ tubes (ThermoFisher Scientific) and incubated for 5 minutes. 100 µL chloroform was then added to the tubes, incubated for 3 minutes and then centrifuged for 15 minutes at 12,000 × g at 4°C. The upper aqueous phase was then transferred to RNase-free tubes and a 1:1 volume of 100% ethanol added and mixed. RNA Clean and Concentrator-5 (Zymo Research) was then used to complete the RNA extraction as per the manufacturer’s instructions. RNA samples were then stored at -80°C until required for qPCR or Nanostring gene expression analysis. The primers used for qPCR were: MUC5AC (Hs01365616_m1), SCGB1A1 (Hs00171092_m1), RAGE/AGER (Hs00542584_g1), SFTPB (Hs00167036_m1), ACE2 (Hs01085333_m1), TMPRSS2 (Hs00237175_m1), GAPDH (Hs02758991_g1) and all were purchased from ThermoFisher Scientific.

### NanoString gene expression analysis

Gene expression analysis of apical cell RNA was undertaken using the NanoString® Human Host Response panel (785 gene targets), plus a custom panel of 48 lung-specific gene targets, using a NanoString® nCounter® Pro Analysis System. nCounts of mRNA transcripts were normalised using the geometric means of 12 housekeeping genes (*ABCF1, ALAS1, GUSB, HPRT1, MRPS7, NMT1, NRDE2, OAZ1, PGK1, SDHA, STK11IP* and *TBP*). Data were analysed and fold-changes, p-values and global significance pathway analysis scores generated by ROSALIND® with criteria provided by NanoString®, outlined in the nCounter® Advanced Analysis 2.0 User Manual. P-value adjustment for false discovery rates (FDR) was performed using Benjamini-Hochberg procedure. Data were considered significant if there was a greater than a 2-fold change and adjusted p-values were less than 0.05.

### Statistical Analysis

Volcano, and violin plots were generated on GraphPad Prism (Version 10). Median absolute log2 fold changes were compared the Wilcoxon matched-pairs signed-rank test or Friedman test with Dunn’s multiple comparison correction. Coefficient of variance and Log2 fold-change comparison plots were produced in R. qPCR and infection efficiency results were compared by students t-test in GraphPad prism (Version 10).

## Results

### Morphological and lung cell type expression differences between MPS and ‘static’ models - monoculture

To determine the utility of lung MPS models for characterising infections, alveolar and bronchial models were developed under static and dynamic flow conditions and characterised for their ability to recapitulate specific lung phenotypes. Both cultures consisted of the same Transwell®, primary lung epithelium-based format and were cultured and differentiated using the same methods. In the dynamic flow MPS condition, the addition of perfused media was introduced, whereby media is flowed around the basolateral side of the Transwell® in the culture well (Figure 1A).

**Figure 1.**
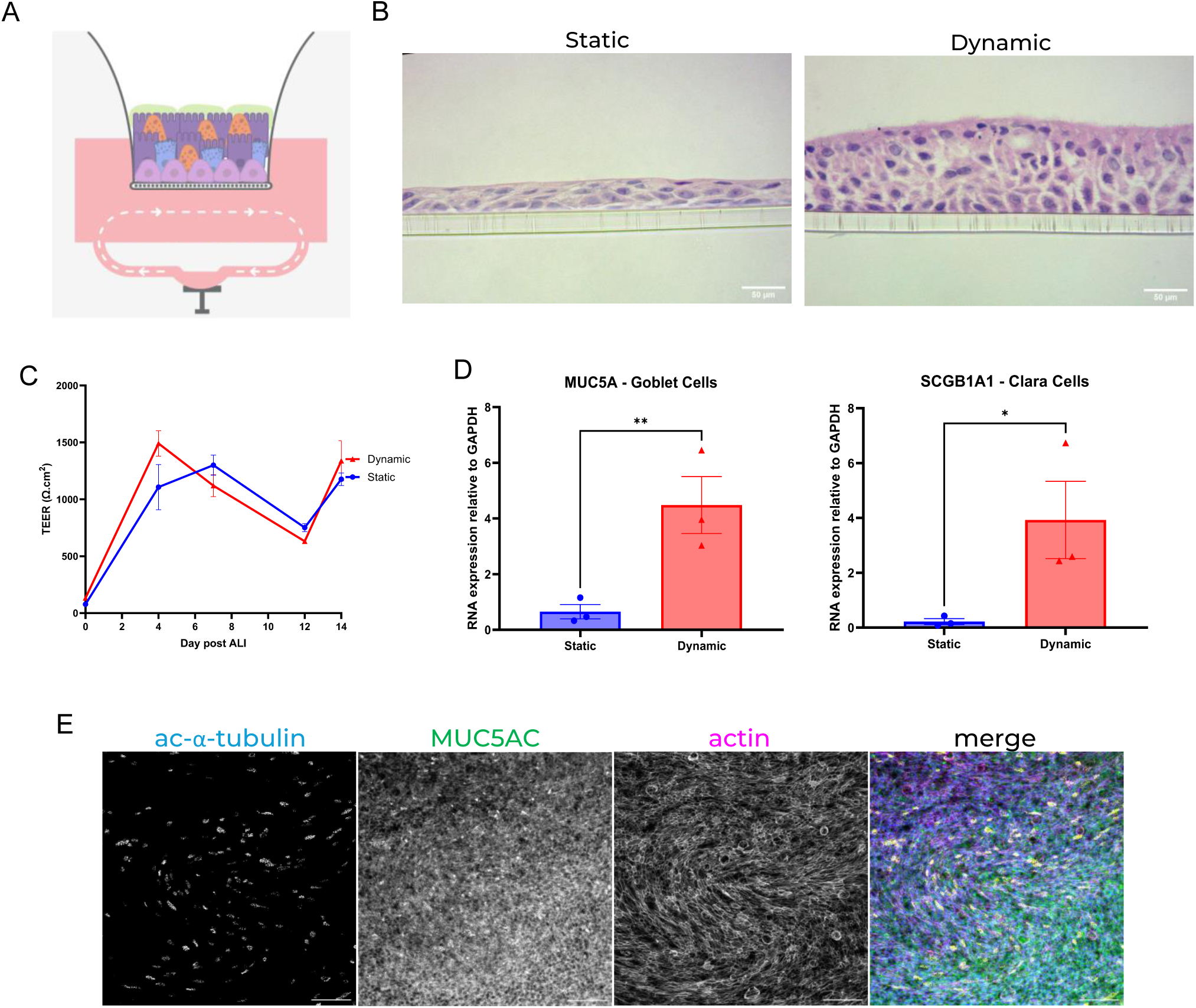
Dynamic flow bronchial MPS promotes enhanced epithelial differentiation compared to static culture. (A) Schematic of the PhysioMimix® bronchial MPS, showing human primary bronchial epithelial cells cultured at air–liquid interface (ALI) under static and dynamic flow conditions. (B) Representative H&E-stained histological sections of bronchial tissues after 14 days of differentiation under static or MPS conditions. Scale bar, 50 µm. (C) Trans-epithelial electrical resistance (TEER) measurements over the 14-day ALI differentiation period. (D) Gene expression of Club cell (SCGB1A1) and Goblet cell (MUC5AC) markers in static versus MPS cultures after 14 days. GP = 0.0332 (*), 0.0021 (**). (E) Immunofluorescence staining of MPS bronchial tissue for acetylated α- tubulin (yellow), mucus (MUC5AC, green), actin (phalloidin, magenta), and nuclei (Hoechst 33342, blue). Scale bar, 100 µm.

Fourteen days after differentiation at air–liquid interface (ALI), tissues were sectioned and analysed for morphological and phenotypic differences. Histological analysis (H&E staining) of NHBE cells cultured under dynamic flow MPS conditions revealed the formation of a pseudostratified epithelium, closely resembling native bronchial tissue (Figure 1B). In contrast, static cultures appeared less developed, despite transepithelial electrical resistance (TEER) measurements showing no significant difference between the two conditions (Figure 1C). qPCR-based analysis of cell composition showed significantly higher levels of Club and Goblet cells in dynamic flow MPS cultures compared to static counterparts (Figure 1D). Immunofluorescence microscopy confirmed the presence of both cilia and mucus in static and dynamic flow cultures (Figure 1E).

Similarly, SAECs were seeded onto Transwell® inserts and differentiated under either static or dynamic flow conditions for 14 days (Figure 2A). Histological analysis demonstrated that tissues exposed to dynamic flow developed complex structures, whereas static cultures remained as uniform, flattened cell layers (Figure 2B). Notably, dynamic flow cultures formed protruding sac-like structures reminiscent of alveoli, creating air pockets throughout the tissue. TEER measurements supported the conclusion that exposure to dynamic flow enhanced tissue structure and differentiation (Figure 2C). Gene expression analysis of key alveolar cell markers - AQP5 (AT1) and SFTPB (AT2) - indicated improved maintenance of both AT1 and AT2 populations in dynamic flow MPS cultures over the 14-day period (Figure 2D). These findings were corroborated by immunofluorescence imaging for RAGE/AGER and SFTPB (Figure 2E).

**Figure 2.**
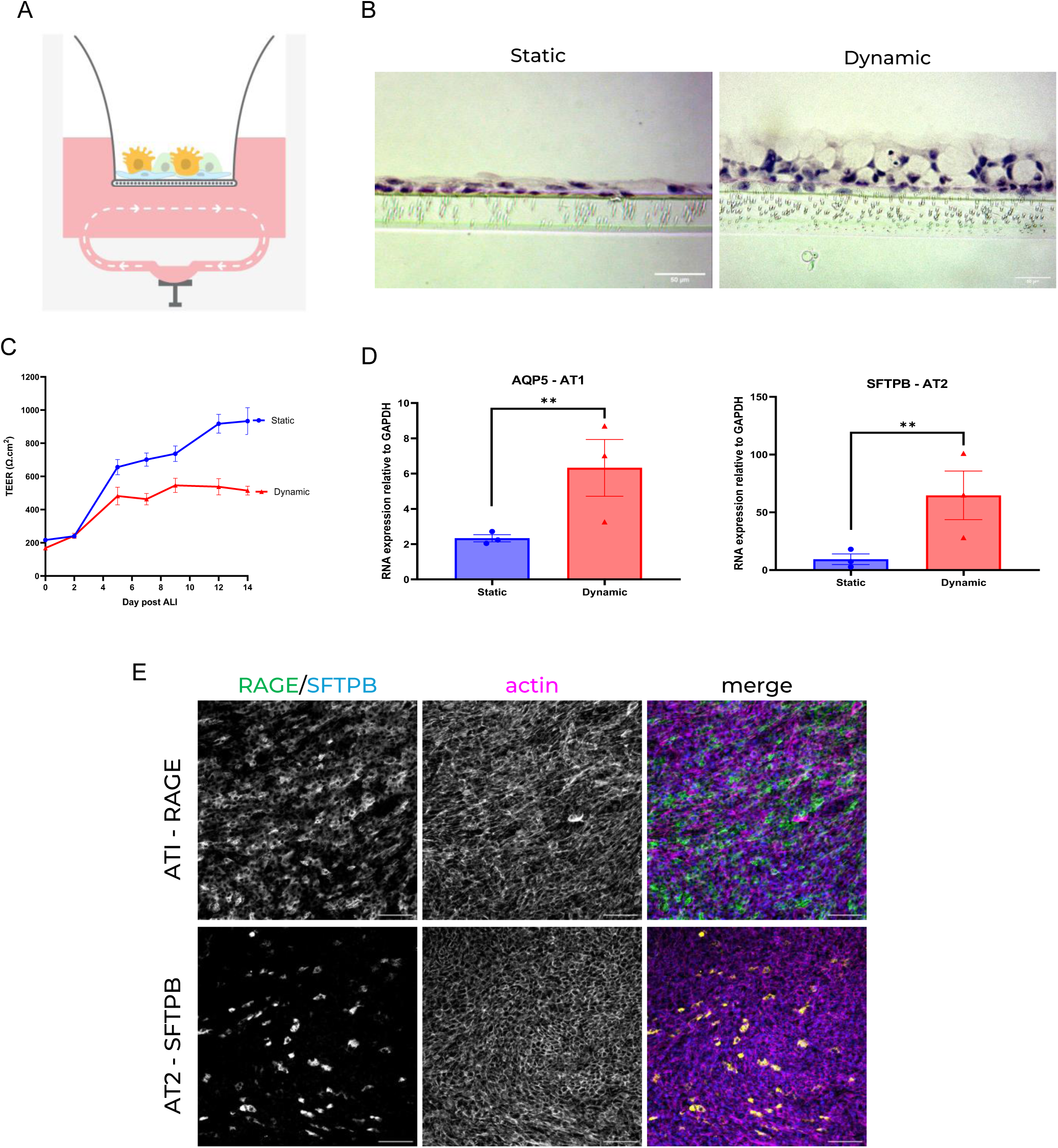
Dynamic flow alveolar MPS promotes enhanced and sustained differentiation into alveolar tissue architecture and phenotype compared to static culture. (A) Schematic of the PhysioMimix® alveolar MPS, showing human primary small airway epithelial cells cultured at air– liquid interface (ALI) under static and dynamic flow conditions. (B) Representative H&E-stained histological sections of alveolar tissues after 14 days of differentiation. Scale bar, 50 µm. (C) Trans-epithelial electrical resistance (TEER) measurements over the 14-day ALI differentiation period. (D) Gene expression of alveolar type I (AT1; AQP5) and alveolar type II (AT2; SFTPB) cell markers in static versus MPS cultures. GP = 0.0021 (**). (E) Immunofluorescence staining of alveolar MPS tissue for AT1 cells (RAGE, green), AT2 cells (SFTPB, yellow), actin (phalloidin, magenta), and nuclei (Hoechst 33342, blue). Scale bar, 100 µm.

### Morphological and lung cell type expression differences between dynamic flow and static cocultures

To further enhance the physiological relevance of the models, primary pulmonary endothelial cells were incorporated on the basolateral side of the Transwell® inserts (Figure 3A). Endothelial cells play a key role at the lung–vascular interface, particularly in mediating inflammatory responses. Upon addition, they formed a thin, uniform monolayer across the underside of the membrane, as shown in histological sections (Figure 3B). While the endothelium alone does not constitute a barrier, its presence increased TEER values in both bronchial and alveolar models, suggesting that epithelial–endothelial interactions enhance epithelial barrier integrity *in vitro* (Figure 3C). However, no significant differences in TEER were observed between dynamic flow and static conditions in either model (Figure 3D). Peak TEER values were comparable between bronchial and alveolar tissues (1859.82 Ω·cm² and 1766.23 Ω·cm², respectively).

**Figure 3.**
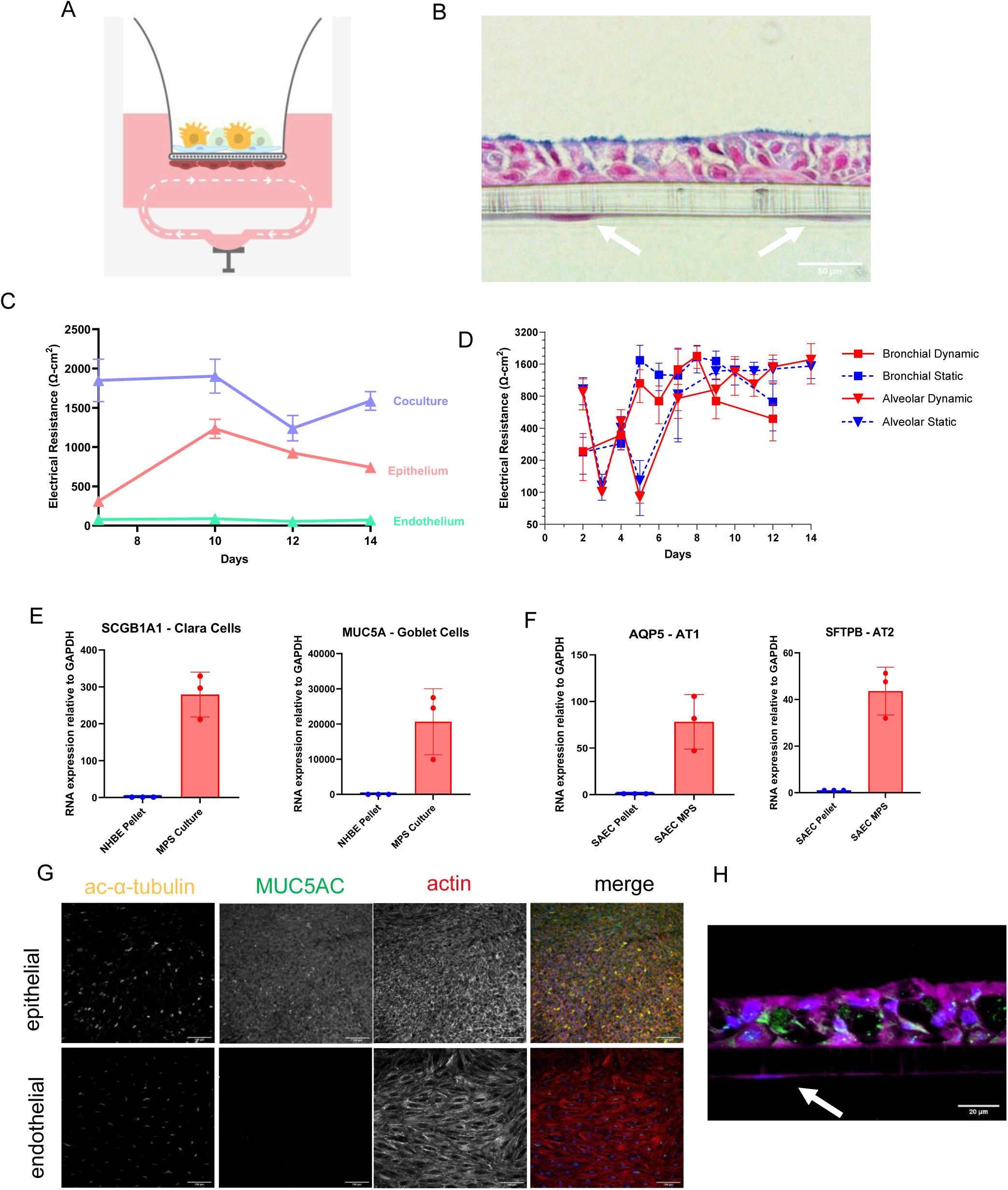
Addition of endothelial cells supports maintenance of epithelial differentiation in lung MPS co-cultures. (A) Schematic of the PhysioMimix® lung MPS co-culture system. Epithelial cells were seeded on the apical side of the Transwell® insert and human pulmonary microvascular endothelial cells (HPMVECs) on the basolateral side, under air–liquid interface (ALI) and dynamic flow conditions. (B) Representative H&E and Alcian blue-stained histological section of bronchial MPS co-culture after 14 days. Mucus is stained blue; endothelial cells are indicated by white arrows. Scale bar, 50 µm. (C) TEER measurements comparing epithelial monoculture, endothelial monoculture, and epithelial–endothelial co-culture over 14 days under ALI conditions. (D) TEER comparison of bronchial and alveolar co-cultures grown under static or dynamic flow MPS conditions over the 14-day ALI differentiation period. (E) Gene expression of Club cell (SCGB1A1) and Goblet cell (MUC5AC) markers in NHBE cells before culture (NHBE pellet) and after 14 days of differentiation in MPS co- culture. (F) Gene expression of alveolar markers - AT1 (AQP5) and AT2 (SFTPB) - in SAEC cells before culture (SAEC pellet) and after 14 days of MPS co-culture. (G) Immunofluorescence staining of bronchial MPS co-culture tissue for acetylated α-tubulin (yellow), mucus (MUC5AC, green), actin (phalloidin, red), and nuclei (Hoechst 33342, blue). Top row shows epithelial layer; bottom row shows endothelial layer. Scale bar, 100 µm. (H) Immunofluorescence staining of alveolar MPS co-culture tissue after 14 days of differentiation, showing surfactant (SFTPB, green), actin (phalloidin, magenta), and nuclei (Hoechst 33342, blue). Endothelial cells are marked with white arrows. Scale bar, 20 µm.

Confocal microscopy and immunohistochemistry confirmed the formation of a consistent endothelial monolayer in both bronchial and alveolar models (Figure 3G–H). Notably, the presence of endothelial cells supported the maintenance of differentiated epithelial phenotypes in dynamic flow MPS cultures of both tissue types (Figure 3E–F).

### Gene expression analysis shows increased separation between bronchial and alveolar models in dynamic flow MPS, compared to static tissues

RNA was extracted from cell lysates collected 7 days post-infection and compared with uninfected controls cultured in parallel. Gene expression was analysed using the NanoString Host Response panel alongside a custom lung-specific panel. When comparing static and dynamic bronchial tissues there were no observable differences in gene expression. Similarly, no significant gene expression differences were observed between static and dynamic alveolar tissues (Supplementary File 2). However, comparison between bronchial and alveolar models revealed 23 differentially expressed, lung region– specific genes (Figure 4 and Table 1).

**Figure 4.**
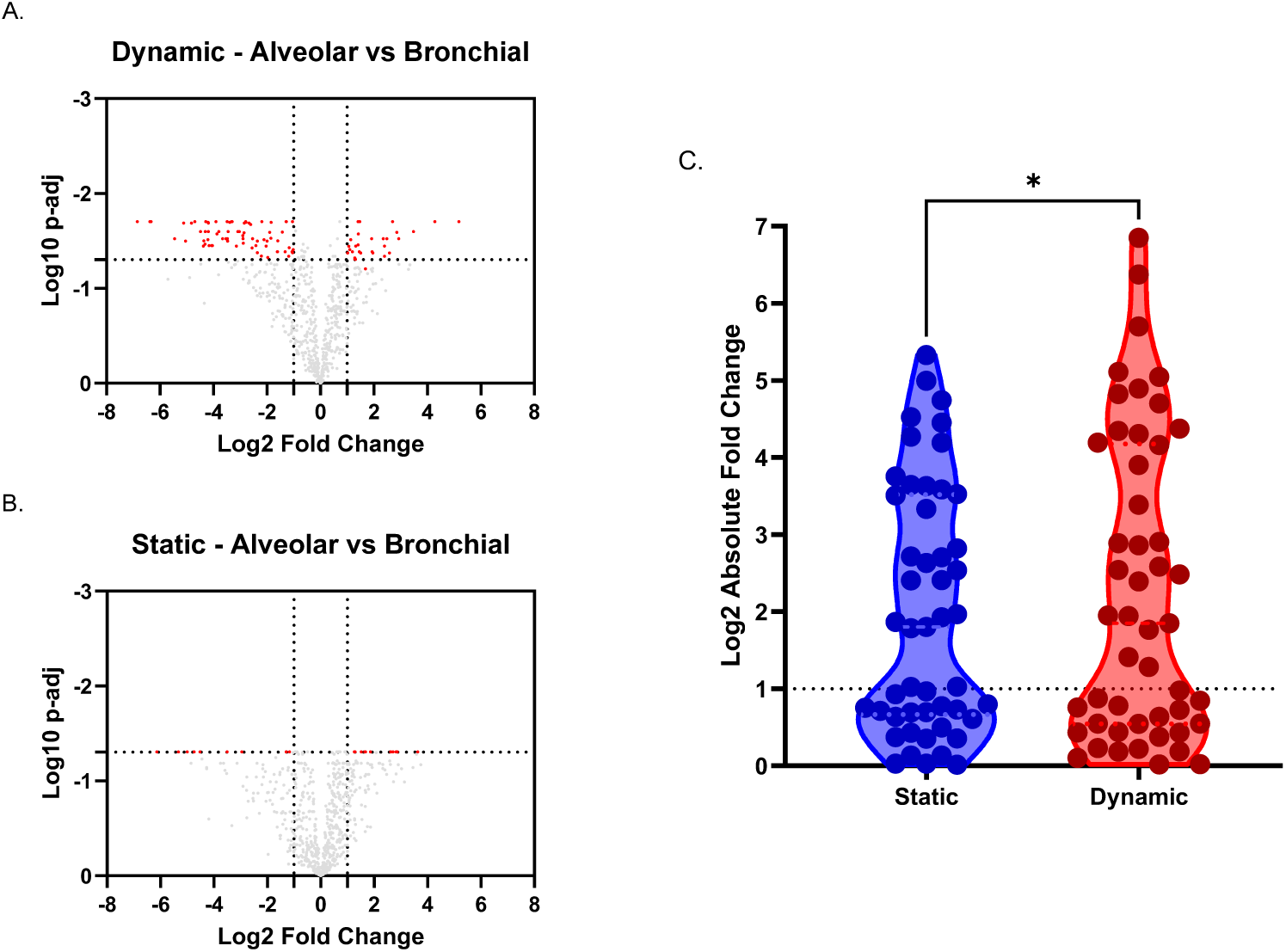
Differential expression of lung-specific and host response genes in alveolar and bronchial models under static and dynamic flow MPS conditions. (A–B) Volcano plots showing differential gene expression from the NanoString Host Response panel and 48 lung-specific gene targets, comparing alveolar and bronchial models cultured under dynamic flow (A) and static (B) conditions. P-values were calculated in ROSALIND® using NanoString guidelines, and adjusted p-values (p-adj) were generated using the Benjamini–Hochberg method to estimate false discovery rate (FDR). Unadjusted p-values are shown in blue; FDR-adjusted p-values are shown in red. Horizontal dotted line indicates p = 0.05; vertical dotted lines indicate ±2-fold change. (C) Violin plot showing absolute log₂ fold changes in lung- specific gene expression between alveolar and bronchial tissues under static and MPS conditions. Dotted line represents a 2-fold change threshold. Statistical significance was assessed using a Wilcoxon signed-rank test; * p < 0.05. Data represent n = 4 lung-on-a-chip models across three independent experiments.

**Table 1.** List of top differentially expressed lung-specific genes identified by NanoString analysis when comparing alveolar to bronchial airway models.

|  | Static |  | Dynamic |  |
| --- | --- | --- | --- | --- |
| Gene | Fold Change | p-adj | Fold Change | p-adj |
| MSMB | -31.923 | 0.049 | -115.298 | 0.020 |
| SERPINB3 | -40.229 | 0.049 | -83.014 | 0.020 |
| MUC5B | -21.889 | 0.059 | -34.563 | 0.020 |
| TMEM190 | -19.294 | 0.074 | -33.000 | 0.032 |
| BPIFB1 | -6.565 | 0.163 | -28.301 | 0.020 |
| UPK1B | -23.007 | 0.049 | -26.012 | 0.020 |
| MKI67 | -26.750 | 0.049 | -20.750 | 0.036 |
| SPDEF | -13.500 | 0.065 | -19.750 | 0.025 |
| LYZ | -12.500 | 0.065 | -18.272 | 0.025 |
| DNAAF1 | -11.517 | 0.083 | -17.917 | 0.035 |
| S100A8 | -11.389 | 0.049 | -14.961 | 0.020 |
| MUC5AC | -5.317 | 0.091 | -10.469 | 0.020 |
| SCGB1A1 | -2.034 | 0.475 | -7.497 | 0.028 |
| SLC26A4 | -3.443 | 0.132 | -5.821 | 0.030 |
| PIGR | -1.710 | 0.404 | -3.603 | 0.030 |
| SLC12A2 | -1.525 | 0.079 | -1.554 | 0.048 |
| SFTPB | 12.385 | 0.049 | -1.463 | 0.731 |
| HOPX | 3.643 | 0.049 | -1.137 | 0.818 |
| RARRES1 | 6.191 | 0.049 | 3.865 | 0.061 |
| CFTR | 6.526 | 0.049 | 5.595 | 0.030 |
| EGLN3 | 5.301 | 0.077 | 6.002 | 0.042 |
| NREP | 7.073 | 0.049 | 7.259 | 0.026 |

Bronchial markers such as MSMB (goblet cells) and UPK1B (club cells) were more highly expressed in bronchial models compared to alveolar models under both static and dynamic conditions. Notably, other bronchial-specific markers - including MUC5B, BFIB1, MUC5AC, and SCGB1A1 - were significantly upregulated in dynamic flow bronchial MPS tissues compared to dynamic alveolar MPS, but this distinction was not observed in the static cultures.

Alveolar markers such as *CFTR* and *NREP* were significantly upregulated in alveolar tissues under both static and dynamic flow conditions. Overall, a greater number significantly differentially expressed genes (DEGs) were detected under dynamic flow, primarily driven by the magnitude of fold-changes in dynamic flow MPS compared to static MPS (Figure 4C).

### Pulmonary dynamic flow MPS tissues demonstrate physiologically relevant responses to stimuli and express SARS-CoV-2 entry receptors

To evaluate the immunological responsiveness of the alveolar and bronchial models, toll-like receptor (TLR) agonists - lipopolysaccharide (LPS, TLR4 agonist) and polyinosinic:polycytidylic acid [poly(I:C), TLR3 agonist] - were applied to the apical (epithelial) surface. Cytokine release was measured from the basolateral compartment over a 48-hour period.

Within six hours of stimulation, both bronchial and alveolar dynamic flow MPS models mounted a detectable inflammatory response to poly(I:C), which was sustained over the 48-hour period (Figure 5A). Notably, alveolar tissues secreted higher levels of IP-10 than bronchial tissues. In response to LPS, only the alveolar dynamic flow MPS model showed increased IP-10 expression, consistent with the known low expression of TLR4 in bronchial epithelium.

**Figure 5.**
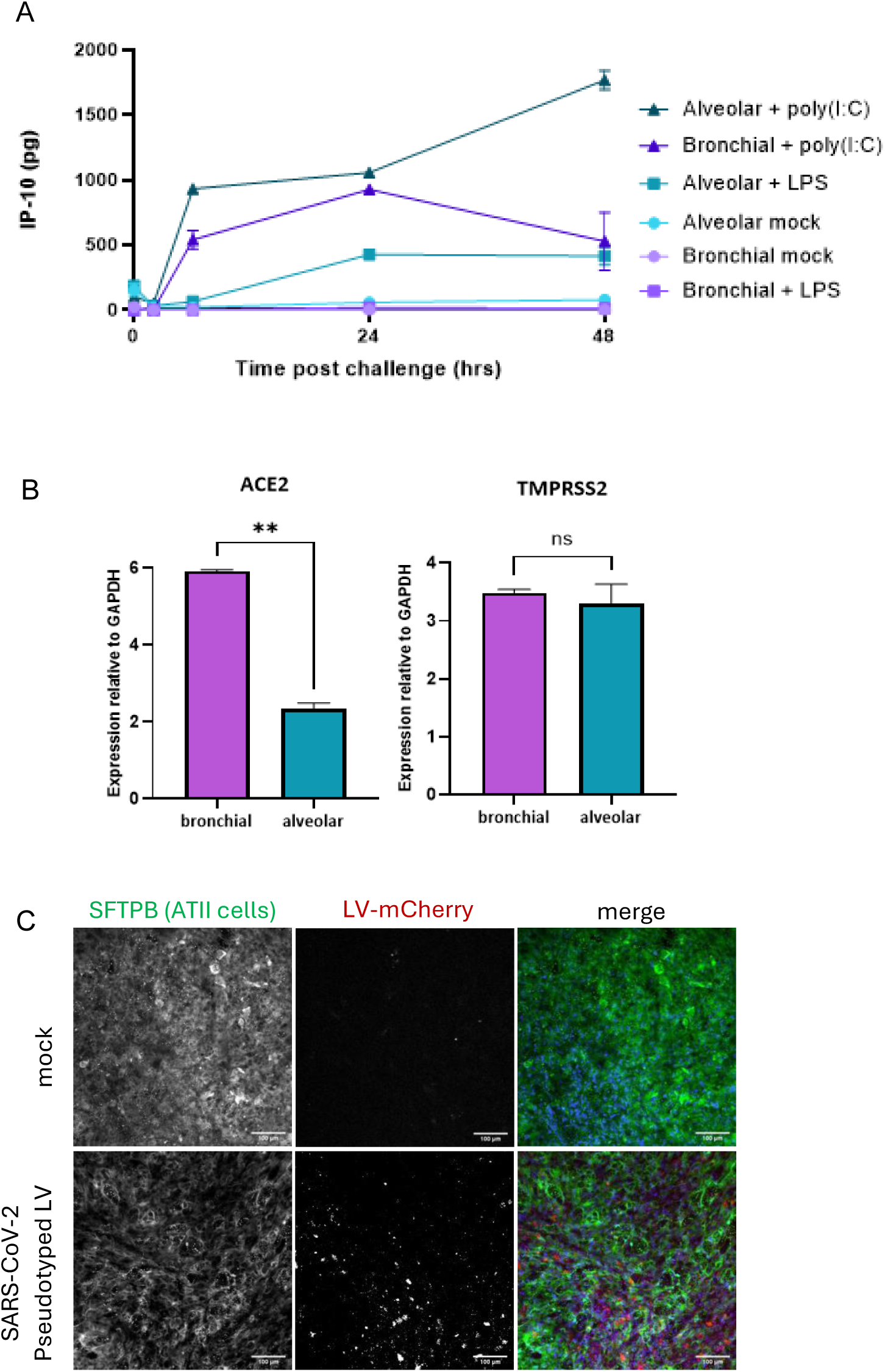
Pulmonary MPS tissues exhibit physiologically relevant inflammatory responses and support pseudotyped SARS-CoV-2 infection. (A) Bronchial and alveolar MPS co-cultures were differentiated for 14 days at air–liquid interface (ALI) under dynamic flow conditions and then challenged with either LPS or poly(I:C). Basolateral supernatants were collected at 2, 6, 24, and 48 hours post-stimulation and analysed for IP-10 secretion. (B) Expression of SARS-CoV-2 entry receptors **ACE2** and **TMPRSS2** in bronchial (purple) and alveolar (blue) MPS tissues following 14 days of ALI culture, quantified by qPCR. (C) Alveolar MPS cultures were infected with SARS-CoV-2 pseudotyped lentivirus (LV) expressing mCherry (red) after 14 days of ALI differentiation. Tissues were fixed after 48 hours and stained for surfactant protein B (SFTPB, green) and nuclei (Hoechst 33342, blue). Scale bar, 100 µm.

To determine the relevance of these models for SARS-CoV-2 research, expression of the viral entry receptors ACE2 and TMPRSS2 was analysed by qPCR (Figure 5B). Both bronchial and alveolar dynamic flow MPS models expressed the receptors, although ACE2 levels were lower in alveolar tissues. TMPRSS2 expression was comparable between the two models. Functional permissiveness to viral entry was confirmed using a SARS-CoV-2 D614G pseudotyped lentivirus expressing EGFP, with infection visualised by immunofluorescence microscopy (Figure 5C).

### SARS-CoV-2 variants infection rate in coculture bronchial and alveolar airway dynamic flow MPS and ‘static’ models

For both static and dynamic flow MPS bronchial airway models, there was no significant difference in infection rate of any of the SARS-CoV-2 variants at an MOI of 1 or 0.01.

At an MOI of 1, infection was successfully established in both bronchial and alveolar airway tissues across all SARS-CoV-2 variants tested (Pre-alpha, Delta, and Omicron BA.5), in both static and dynamic flow MPS conditions (Figure 6A–B). In bronchial models, infection rates were high regardless of culture condition, and no significant differences were observed between static and MPS systems for any variant. However, static bronchial tissues infected with Pre-alpha showed a non-significant reduction in infection rate (67%) relative to Delta and Omicron, which both reached 100%. In contrast, alveolar models exhibited greater variability in infection efficiency. Omicron achieved 100% infection across both systems, whereas Pre-alpha infection rates were significantly lower: 47% in dynamic flow MPS (p = 0.003) and 43% in static (p = 0.009). Delta infection in alveolar tissues was also reduced in static cultures (43%) relative to Omicron (p = 0.009), with a non-significant trend toward reduced infection in MPS models (69%, p = 0.052). No significant difference in infection rate was detected between static and MPS alveolar tissues for any individual variant.

**Figure 6.**
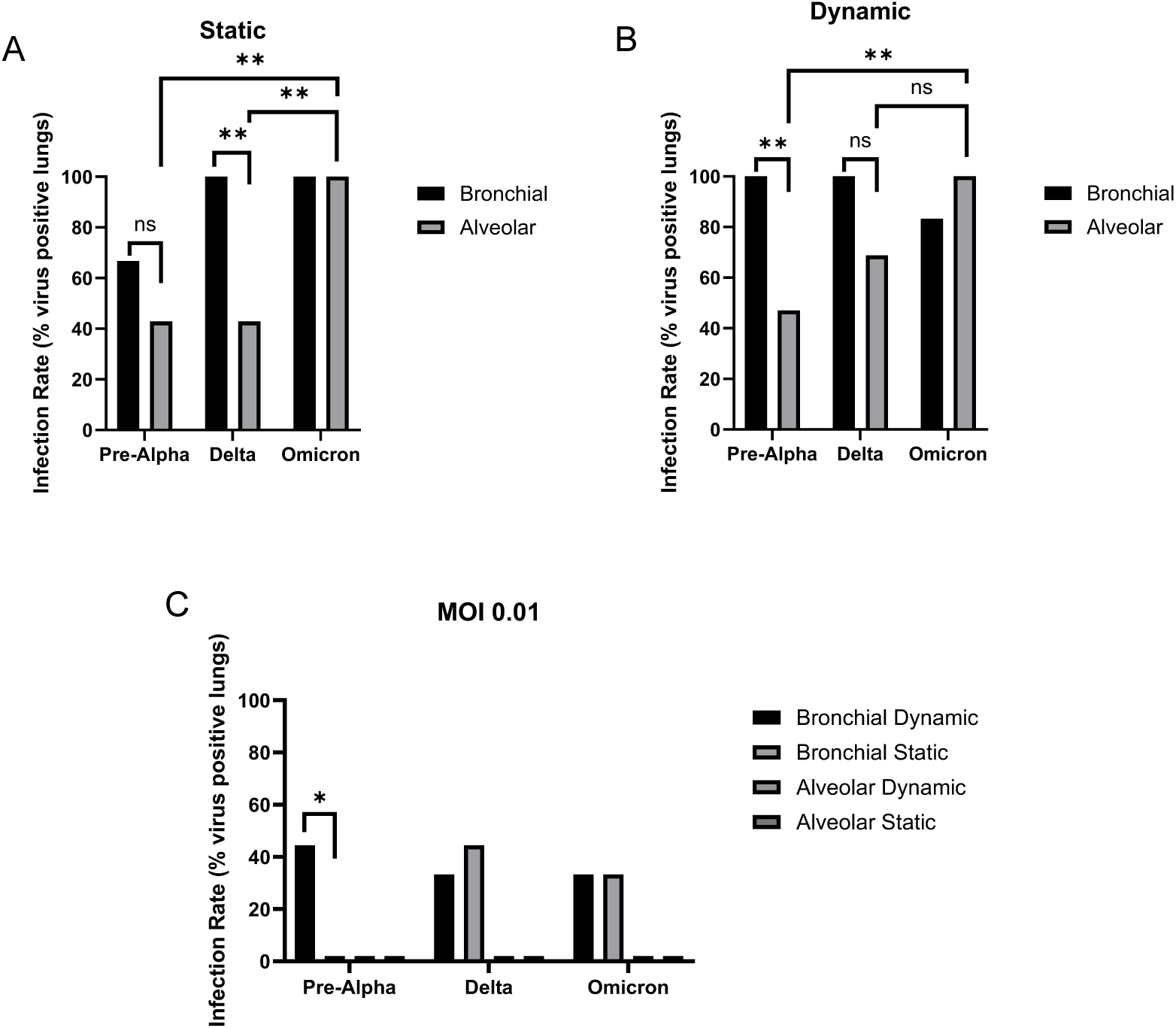
Infection efficiency of SARS-CoV-2 in static and dynamic flow MPS bronchial and alveolar airway models. Lung tissues cultured in static or dynamic flow MPS conditions were infected with SARS-CoV-2 variants pre-alpha, delta and omicron at an MOI of 1 (A and B) or 0.01 (C). Apical surface washes were collected on day 2, 4 and 7 post-infection and live SARS-CoV-2 was quantified by VeroE6 plaque assay. All plaque assays were conducted with three technical replicates. lung tissues were classed as having a successful infection if any live SARS-CoV-2 was recovered on day 2, 4 or 7 post-SARS-CoV-2 infection. Data shown is from n = 4-17 lung tissues, from 3 independent experiments and was analysed by two-tailed t-test, ns = not significant ** = p<0.01, * = p<0.05

At an MOI of 0.01, infection rates were markedly reduced across all conditions (Figure 6C). In bronchial tissues, Pre-alpha infection was significantly higher in dynamic flow MPS (44%) compared to static (0%, p = 0.022), while Delta and Omicron showed similar infection rates between conditions, with no significant differences. Alveolar tissues did not support infection at this lower MOI in either static or MPS cultures. Based on these initial findings, an MOI of 1 was selected for all subsequent experiments, as it reliably established infection in both bronchial and alveolar airway models under static and dynamic flow conditions.

### SARS-CoV-2 variant infection dynamics

To assess infection dynamics, apical surface washes were collected at 0, 2, 4, and 7 days post-infection. Viral replication was quantified by qPCR and plaque assay using VeroE6 cells. Across all SARS-CoV- 2 variants, peak viral titres were observed at day 4, with levels either plateauing or declining by day 7 (Figure 7). Differences in infection dynamics were evident between bronchial and alveolar models. For Pre-alpha, bronchial tissues exhibited a 10- to 100-fold higher viral titre compared to alveolar tissues. Delta (B.1.617.2) infection also showed elevated replication in bronchial tissues, though to a lesser extent (6- to 10-fold increase). In contrast, Omicron (BA.5) infection resulted in similar levels of viral replication across both airway models (Figure 7). This pattern was mirrored in qPCR quantification of viral RNA, although the magnitude of the observed differences was lower (Figure 7).

**Figure 7.**
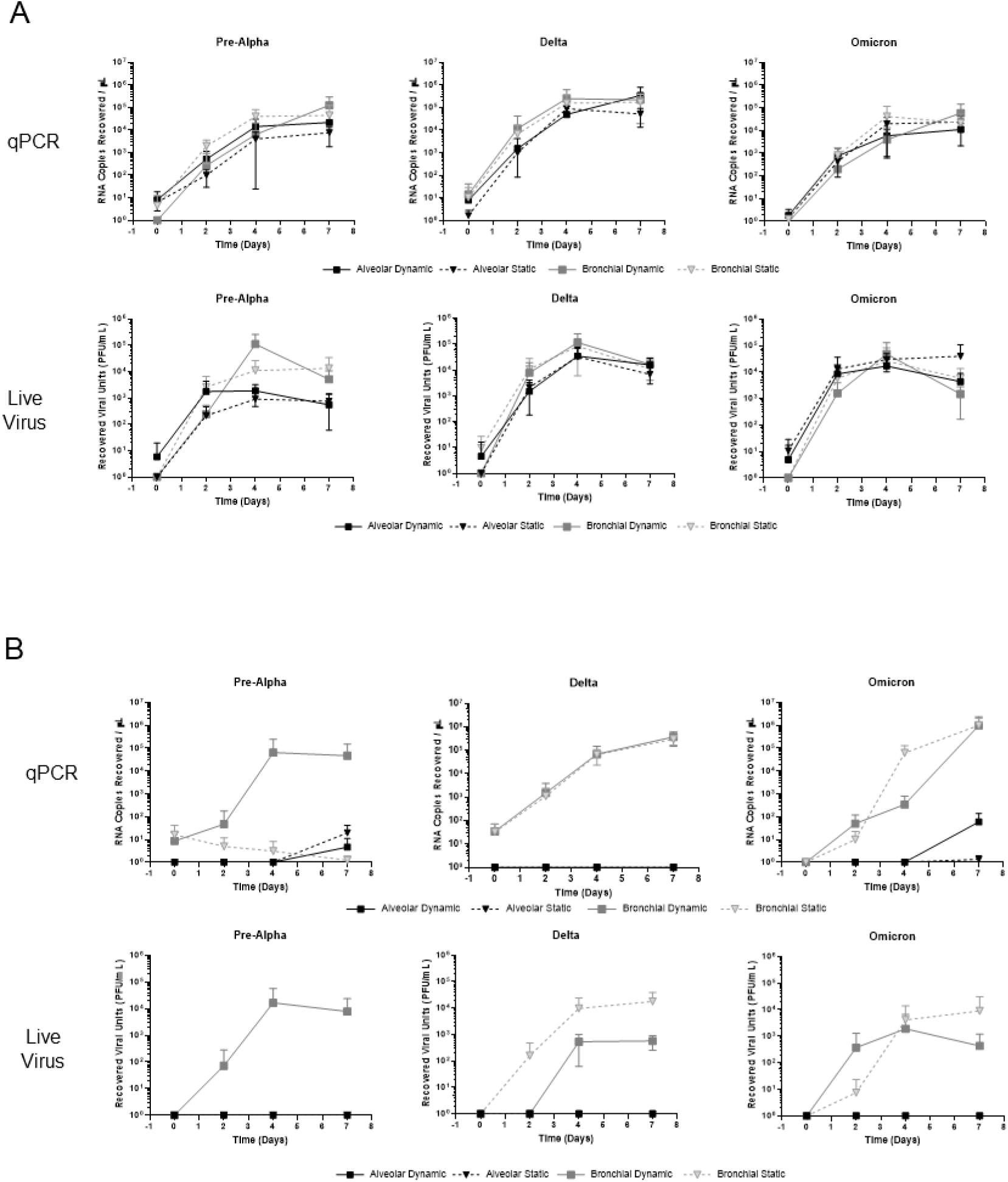
Infection dynamics of bronchial and alveolar airway models following SARS-CoV-2 infection at MOI 1 or 0.01. Bronchial and alveolar airway tissues were infected with SARS-CoV-2 variants - Pre-alpha (2020 reference strain), Delta (B.1.617.2), and Omicron (BA.5) - at a multiplicity of infection (MOI) of 1 (A) or 0.01 (B) for two hours. After infection, viral inoculum was removed, and tissues were washed three times with PBS before incubation under air–liquid interface (ALI) conditions for 7 days. Apical washes were collected on days 0, 2, 4, and 7 post-infection. Viral RNA was quantified by qPCR, and infectious virus present in apical washes was measured by plaque assay using VeroE6 cells to assess infection dynamics across airway regions and viral variants. Data represent n = 4–17 lung tissues from three independent experiments. Plaque assays were performed in technical triplicate.

At seven days post-infection, tissues were fixed and stained for SARS-CoV-2 nucleocapsid protein (Abcam, ab280201), followed by detection with Alexa Fluor® 488-conjugated secondary antibody (Goat Anti-Rabbit IgG H&L). Fluorescence microscopy revealed variant-specific infection patterns.

In bronchial tissues, distinct differences in staining were observed between variants. Delta (B.1.617.2) showed widespread infection, characterised by large syncytia formation and signs of nuclear DNA damage, indicated by intense, punctate nuclear staining (Figure 8). In contrast, Omicron (BA.5) infection was localised to discrete, concentrated foci in both bronchial and alveolar tissues (Figure 8), a pattern less frequently observed in Pre-alpha and Delta infections. Overall, alveolar models showed smaller, more confined regions of infection and reduced syncytia formation compared to bronchial tissues.

**Figure 8.**
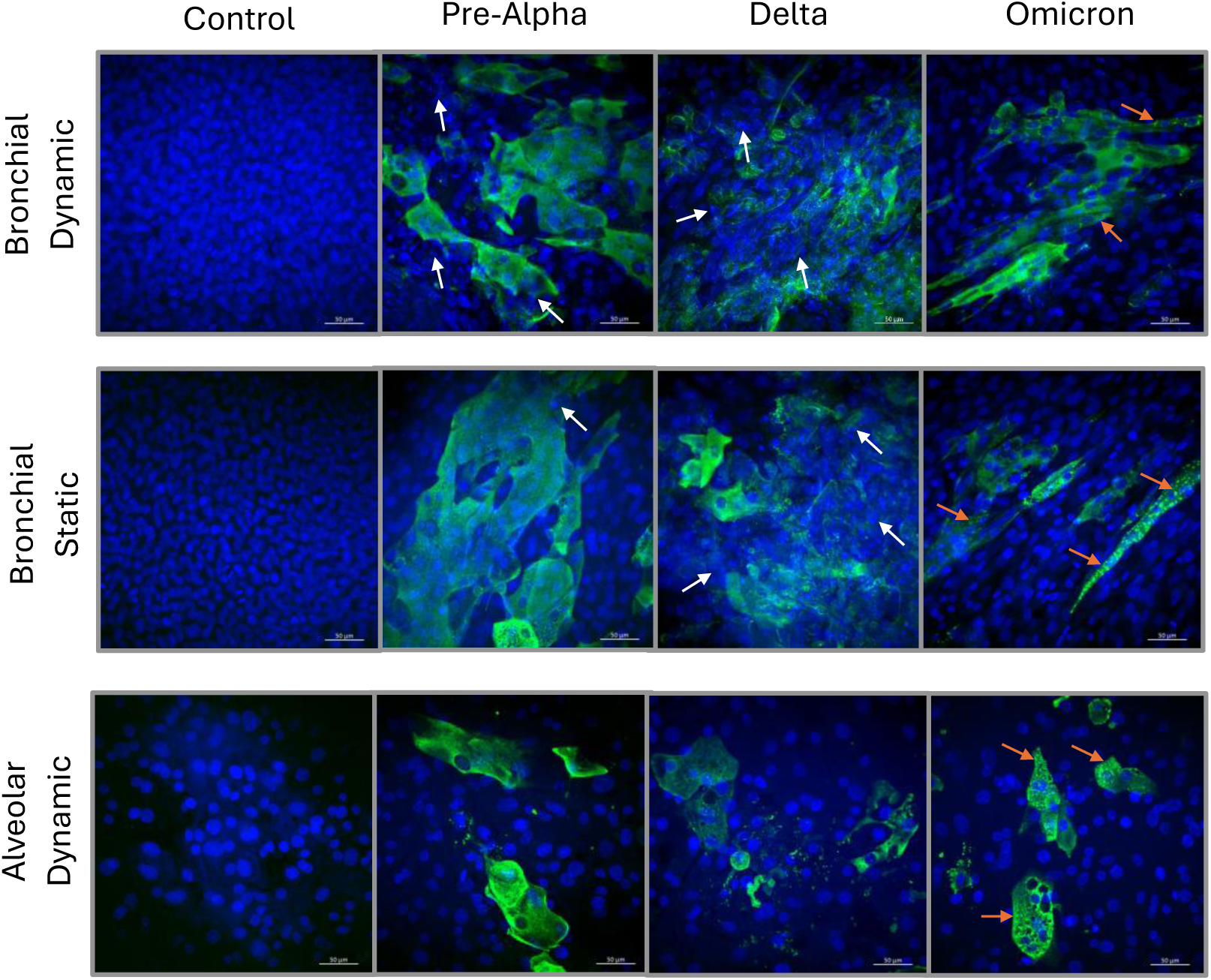
SARS-CoV-2 displays variant specific morphology of infection in static dynamic flow MPS bronchial and alveolar tissues. Static and dynamic flow MPS bronchial and alveolar tissues were infected with the SARS-CoV-2 variants, pre-alpha (2020 early reference strain), delta B.1.617.2 and omicron BA.5 at an MOI of 1 for two hours. After which, SARS-CoV-2 containing media was removed and lung tissues were washed three times with PBS. lung tissues were incubated for 7 days and then fixed with 4% formaldehyde and stained with Anti-SARS-CoV-2 nucleocapsid protein antibody [EPR24334-118](Abcam, ab280201), followed by Goat Anti-Rabbit IgG H&L, Alexa Fluor® 488 (Abcam, ab150077) (shown in green) and DAPI (shown in blue). DNA damage indicated by white arrows and omicron high intensity fluorescence pockets shown with orange arrows. Lung tissues membranes were mounted onto slides and imaged using Zeiss confocal microscope at 40x objective. Representative images are shown from n = 3 lung tissues from three separate experiments.

### Differences in the number of DEGs are observed between models and SARS-CoV-2 variants, but interferon response pathways are activated regardless of variant

Cell lysates were collected 7 days post-infection, and RNA was extracted for gene expression analysis using the NanoString Host Response Panel alongside a custom lung-specific panel. Across all models, infection with Delta (B.1.617.2) resulted in the highest number of DEGs with an FDR-adjusted p-value ≤ 0.05 (Figure 9A). In bronchial tissues, Pre-alpha induced the second-highest number of DEGs, followed by Omicron (BA.5). In contrast, this pattern was reversed in alveolar tissues, where Omicron induced more DEGs than Pre-alpha. Delta infection consistently resulted in larger overall fold-changes compared to Pre-alpha or Omicron across both models, as assessed by Friedman test with Dunn’s correction for multiple comparisons (Figure 9B-D; p < 0.0001).

**Figure 9.**
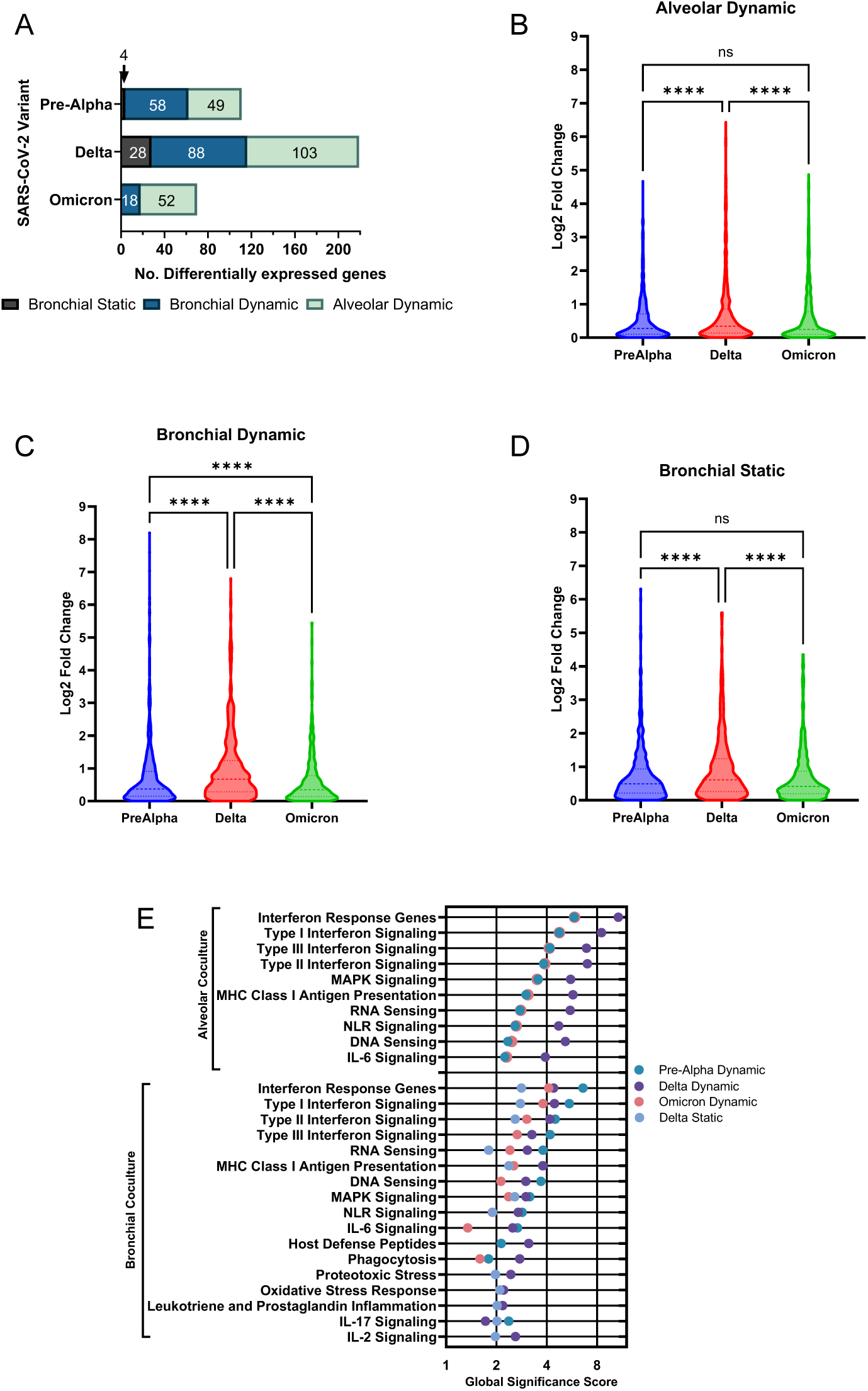
Number of significantly differential expressed host response and lung panel genes, absolute fold change gene expression and top signalling pathways influenced following SARS- CoV-2 variant infection in bronchial and alveolar tissues. Dynamic flow MPS and static tissues were infected with SARS-CoV-2 variants for 7 days. At the experimental endpoint, apical cells were collected, RNA extracted, and gene expression quantified with NanoString Host Response panel and custom lung panel. (A) The number of statistically significant differentially expressed gene changes, defined as FDR adjusted p-value ≤0.05 with a fold-change greater than 2, compared to uninfected controls, are displayed for pre-alpha, delta and omicron infected static bronchial tissue (black), dynamic flow MPS bronchial tissue (blue) and dynamic flow MPS alveolar tissue (green). Median log2 absolute fold-change values of Pre-Alpha, Delta and Omicron differentially expressed genes in dynamic flow MPS alveolar (B) dynamic flow MPS bronchial (C) and static bronchial (D) tissues were compared by Friedman test with Dunn’s correction for multiple tests. ns = not significant, **** = p<0.0001. (E) Significantly differentially expressed genes identified by ROSALIND software analysis and global significance scores were generated for host response panel signalling pathways. The pathways with the largest global significance scores in alveolar and bronchial tissues following infection with Pre-Alpha, Delta or Omicron for 7 days are shown. Global significance scores were not generated for static Pre- Alpha and Omicron bronchial infected cocultures as too few genes were significantly differentially expressed.

A greater number of DEGs were observed in bronchial dynamic flow MPS cultures compared to their static counterparts (FDR-adjusted p ≤ 0.05; Figure 9A). To determine whether this was driven by larger effect sizes or reduced variability, log₂ fold change was plotted against the coefficient of variation (Figure 10). The analysis revealed that the increase in DEGs in dynamic flow MPS was attributable to both greater fold-changes and reduced variability across replicates. This effect was particularly evident in bronchial tissues infected with the Pre-alpha variant.

**Figure 10.**
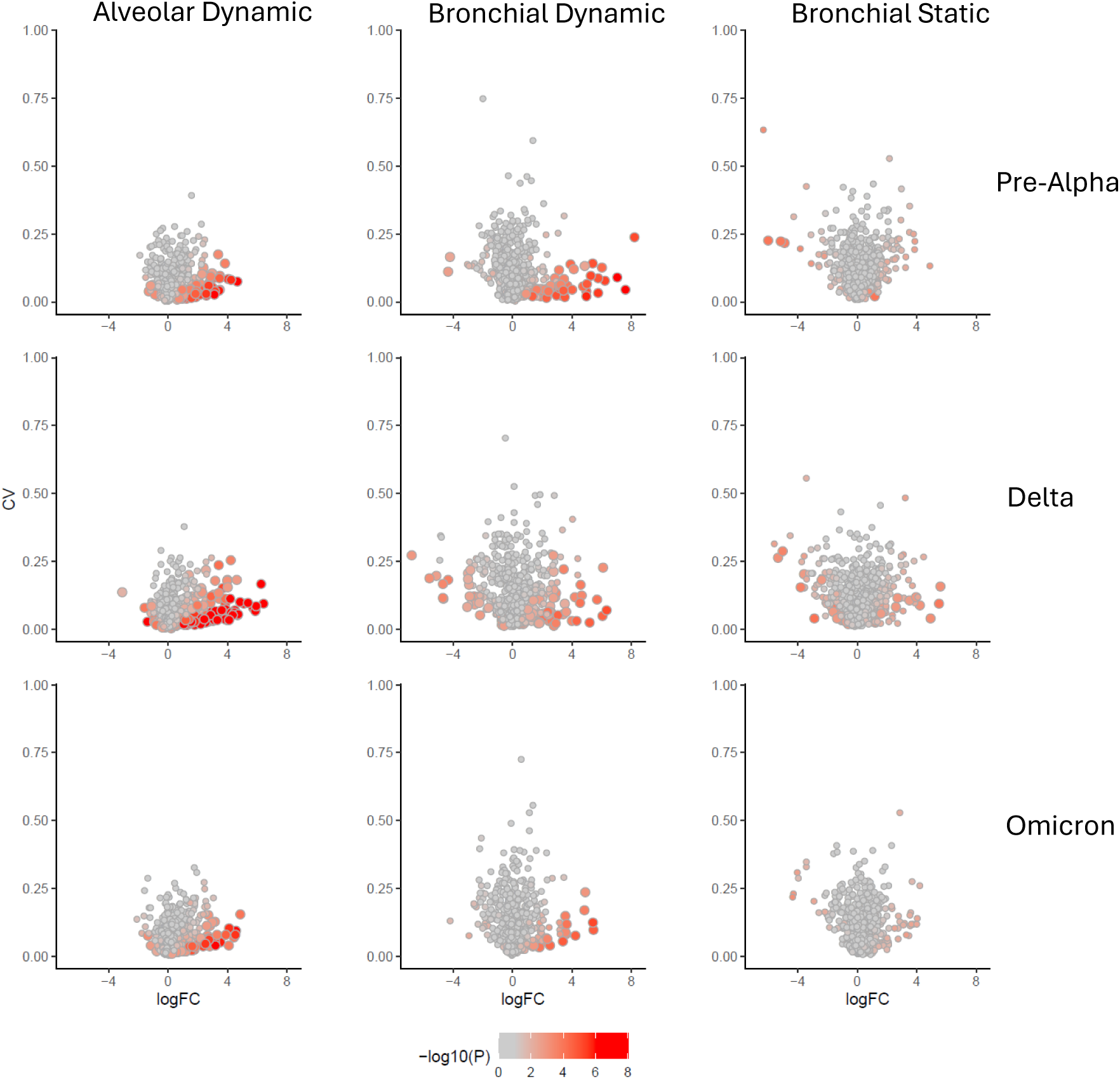
Differential expression and variability of host response and lung-specific genes in bronchial and alveolar tissues following SARS-CoV-2 infection. Differentially expressed genes from NanoString Host Response and lung-specific panels are plotted as log₂ fold change (logFC) versus coefficient of variation (CV). Genes with p-values > 0.05 are shown in grey, while those with p ≤ 0.05 are coloured in shades of red, as indicated in the figure legend. Genes with FDR-adjusted p-values ≤ 0.05 are highlighted with larger circles. Columns indicate the tissue model (static or dynamic flow MPS), and rows indicate the SARS-CoV-2 variant used for infection.

Gene set enrichment analysis was performed using ROSALIND software to calculate global significance scores for pathway activation. Interferon signalling - specifically type I, II, and III pathways - emerged as the most significantly enriched response across all models and variants (Figure 9E). These pathways were consistently activated in alveolar tissues across all variants and were largely preserved in bronchial models. However, additional pathway activations specific to Delta infection were identified, including proteotoxic stress, oxidative stress, leukotriene and prostaglandin signalling, and IL-2 signalling (Figure 9E).

Global significance scores could not be generated for static bronchial tissues infected with Pre-alpha or Omicron due to the low number (or absence) of DEGs meeting the FDR ≤ 0.05 and log₂ fold-change > 1 threshold.

## Discussion

In infectious disease research, a high attrition rate of therapeutics and vaccine candidates at the clinical testing stage is thought to be driven, in part, by the lack of *in vitro* and *in vivo* animal models that accurately replicate the site of infection [11]. This became evident during the COVID-19 pandemic where pre-clinical data was generated from inferior *in vitro* models (e.g. VeroE6 and Calu-3 cell models), which were not fully representative. This led to false identification of numerous SARS-CoV- 2 drug candidates which were interpreted as having promising pre-clinical data, but failed when assessed clinically e.g. chloroquine [12].[13]. Several studies have indicated that dynamic flow MPS are better able to screen pre-clinical drug candidates and identify therapeutics which would fail in later clinical trials [8, 14]. However, there have been few studies which have directly compared dynamic flow MPS with static culture systems when investigating fundamental research questions [4, 15] and none, to our knowledge, that have assessed their appropriateness for infectious disease research. We therefore conducted studies directly comparing tissue architecture, lung cell composition and SARS- CoV-2 infection in bronchial and alveolar models grown either with or without dynamic flow MPS.

Here, we have demonstrated that bronchial and alveolar monocultures grown in dynamic flow MPS generate tissues which better resemble human physiology, displaying more clinically comparable morphology and architecture [16, 17] and higher expression of key bronchial and alveolar cell markers, compared to static cultures (Figures 1 and 2). We further show that addition of endothelial cocultures supports retention of the differentiated phenotype - producing mucus and surfactant typical of human bronchial and alveolar airways [18](Figure 3).

NanoString gene expression analysis revealed no significant gene changes between bronchial dynamic flow MPS and static tissues or alveolar dynamic flow MPS and static tissues, as reported elsewhere [15]. It has previously been demonstrated that neither dynamic flow nor mechanical stretching resulted in major differences in gene expression, although media flow did enhance mucociliary maturation relative to static models [2]. This suggests that differences between static and dynamic flow MPS models are more likely to reflect functional differences rather than changes in cell population composition

It is still not fully understood how dynamic flow of media results in morphological changes, however, previous studies have demonstrated that shear-stress in bronchial epithelial cultures led to improved barrier function [19] and regulates airway surface liquid [20, 21] and mucus secretion [22]. We hypothesise that shear stress from dynamic flow induces mechano-transduction signalling within endothelial cells on the basolateral side, triggering crosstalk with epithelial cells and supporting differentiation and the formation of more physiologically relevant structures. We further propose that dynamic flow enhances physiological relevance by promoting more uniform distribution of oxygen and essential nutrients, reducing the formation of steep media gradients, and improving diffusion to the apical surface of epithelial cells. These improvements in microenvironmental stability are likely to directly impact tissue health, epithelial polarity, and functional maturation. In turn, this can influence multiple aspects of host–pathogen interaction, including receptor expression (e.g. ACE2 and TMPRSS2), cytokine secretion profiles, and epithelial barrier integrity - all of which are critical to the outcomes measured in our models. These factors may contribute to the enhanced infection efficiency seen in alveolar tissues, which are otherwise poorly permissive in static conditions.

Greater differentiation between bronchial and alveolar *in vitro* tissues, which we have attained with dynamic flow MPS tissues (Figure 4), including increased ACE2 expression in bronchial tissues, compared to alveolar tissue (Figure 5), provides more physiologically relevant models, which match clinical observations that ACE2 expression decreases down the respiratory tract [23, 24]. This is necessary for respiratory infection studies to facilitate research into areas such as vaccine development and pathogen entry mechanisms using a more appropriately matched model to the airway region of interest. For example, bronchial tissues are more suitable for studies at the primary sites of infection, whilst alveolar tissues should be utilised to study disease progression and pathogenicity, being the site of acute infections, leading to ARDS.

We have confirmed that dynamic flow bronchial and alveolar MPS respond to poly(I:C) and LPS challenge in a manner consistent with previously published data [25–27] and are also able to be infected with both pseudotyped (MOI 100) and native SARS-CoV-2 (MOI 0.01 and 1) (Figure 5 and 6).

Infection of bronchial tissues with three SARS-CoV-2 variants (Pre-alpha, Delta (B.1.617.2) and Omicron (BA.5)) revealed that all variants were able to infect and replicate with near 100% efficiency (Figure 6). However, infection in alveolar tissue revealed significantly reduced infection rates for Pre-Alpha and Delta (B.1.617.2), particularly in static tissues (Figure 6 and 7) and suggest that dynamic flow MPS tissues have an increased ability to support replication of SARS-CoV-2 variants. We hypothesise the difference in infection rate and dynamics between bronchial and alveolar models, as well as observed difference in replication dynamics between SARS-CoV-2 variants is due to several factors. These may include intrinsic differences in ACE2 and TMPRSS2 expression, tissue architecture and oxygenation, as well as variant-specific tropism and entry pathway preference.

As described previously, alveolar tissues have a significantly reduced ACE2 expression (Figure 5B), compared to bronchial tissues and this is the preferred entry mechanism for earlier strains of SARS-CoV-2, such as Pre-alpha and Delta (B.1.617.2), whereas Omicron (BA.5) has adapted to use TMPRSS2 and endocytic entry mechanisms which can overcome the reduced ACE2 expression [28]. Further to this, alveolar tissues generated a larger IP-10 cytokine response to poly(I:C) challenge which Omicron (BA.5) has the potential to overcome via immune evasion strategies that inhibit interferon production and induction of cytokine storms [28, 29]. These findings partially align with clinical observations - such as Delta’s association with more severe pathology and Omicron’s altered tropism - and suggest dynamic flow MPS may offer additional resolution in modelling clinically relevant infection patterns, particularly in alveolar contexts.

Moreover, NanoString analysis revealed *HSP90AB1* upregulated expression in bronchial tissues and has been shown to be a facilitator in SARS-CoV-2 infection [30, 31]. This, in combination with morphological and cell composition differences, increased ACE2 expression, reduced cytokine response and Omicron (BA.5) adaptation may lead to the improved infection efficiency observed in bronchial tissues and by Omicron (BA.5), however, further studies would be needed to confirm this.

Immunofluorescent imaging of SARS-CoV-2 variants revealed distinct infection patterns across bronchial and alveolar tissues, with observationally, Delta (B.1.617.2) appearing to result in the largest number of syncytia and areas of DNA damage, as indicated by small, intensely fluorescing areas of DAPI stain (Figure 8). Clinically, syncytia formation has been used as a marker of advanced disease following SARS-CoV-2 infection from histological sampling [32, 33], whilst *in vitro* studies have similarly found Delta variants to result in the greatest number of syncytia [34]. Future studies will seek to quantitatively characterise these differences with larger data sets which would allow detailed image analysis to be undertaken and identify individual variants unique infection morphology. Imaging techniques in combination with complex airway models could then be utilised to aid prediction of pathogenicity of future variants of concern and/or as a secondary measure of the effectiveness of therapeutic or vaccine interventions.

Our characterisation of the host response to SARS-CoV-2 within the airway models by NanoString also indicated that infection with Delta (B.1.617.2) consistently resulted in the largest number of statistically significant DEGs, due to larger over-all fold changes compared to Pre-alpha and Omicron (BA.5) (Figure 9A-D). Pathway analysis revealed interferon response genes to be the main driver of SARS-CoV-2 infection, such as *CXCL10 and CXCL11* chemokines and *IFIT1, IFIT2, IFIT3* and *ISG15* interferon genes, regardless of variant in bronchial and alveolar models (Figure 9E), in alignment with previously published patient data [35, 36]. This, in together with the observed increase in the number of syncytia, aligns with clinical data which associated Delta variants with increased risk of hospitalisation and severe outcomes [37–39]. These observations also match *in vivo* animal data [40, 41], highlighting, the utility of these models to predict variant pathogenicity.

When comparing bronchial dynamic flow MPS and static tissues, dynamic flow MPS infected models consistently resulted in more differentially expressed genes being identified, due to a combination of larger fold changes and lower coefficient of variance (Figure 10). Dynamic flow MPS cultures provided greater sensitivity for transcriptomic analyses, with improved separation of infected and mock tissues and more consistent responses across replicates.

The data presented here demonstrate our ability to generate complex alveolar and bronchial models under dynamic flow and static culture conditions. Our data demonstrate that there is little difference in infection dynamics and efficiency of static and dynamic flow bronchial MPS infected with SARS-CoV- 2 variants. Therefore, for studies simply assessing infection dynamics, static bronchial models may be sufficient; however, this observation may be limited to studies focused on SARS-CoV-2 and other pathogens may behave differently and further study would be required. For more discrete analysis, such as gene expression quantification, dynamic flow MPS models may be more appropriate as greater confidence can be gained from the data, and fewer replicates required. This is particularly relevant when working with primary and donor cells which are in limited supply. Studies of SARS-CoV-2 infection in the lower airway also benefit from use of dynamic flow alveolar MPS, as our data revealed reduced infection efficiency of SARS-CoV-2 variants in static alveolar models; except for Omicron (BA.5) which was just as infectious regardless of model. We believe that our data demonstrate that dynamic flow MPS provide an exciting tool for infectious disease research - supporting the generation of more physiologically relevant models with the potential to be utilised in future studies to assess therapeutic potency in models which better represent the targeted tissues. These models also lend themselves to provide insights in how donor diversity affects infection dynamics and progression to disease.

## Acknowledgments

This study was supported by the UK Research and Innovation Strength in Places Fund (SIPF 20197, GAB and SHP), the Liverpool City Region Life Sciences Investment Zone (GAB and SHP) and The Pandemic Institute (GAB and SHP).

## Conflicts of Interest

ER and TK are employed by CN Bio, the suppliers of the PhysioMimix® system used for the study.

## References

1. Phan, T.H., et al., Advanced pathophysiology mimicking lung models for accelerated drug discovery. Biomaterials Research, 2023 Apr 26. 27(1).

2. Nawroth, J.C., et al., Breathing on chip: Dynamic flow and stretch accelerate mucociliary maturation of airway epithelium in vitro. Materials Today Bio, 2023/08/01. 21.

3. Gard, A.L., et al., High-throughput human primary cell-based airway model for evaluating influenza, coronavirus, or other respiratory viruses in vitro. Scientific Reports, 2021 Jul 22. 11(1).

4. Dufva, M., A quantitative meta-analysis comparing cell models in perfused organ on a chip with static cell cultures. Sci Rep, 2023. 13(1): p. 8233.

5. Šuligoj, T., et al., Modelling SARS-CoV-2 infection in a human alveolus microphysiological system. Access Microbiology, 2024/09/11. 6(9).

6. Fisher, C.R., et al., *A High-Throughput,* High-Containment Human Primary Epithelial Airway Organ-on-Chip Platform for SARS-CoV-2 Therapeutic Screening. Cells, 2023 Nov 16. 12(22).

7. Quezada, L.L., et al., Predicting Clinical Outcomes of SARS-CoV-2 Drug Efficacy with a High-Throughput Human Airway Microphysiological System. Advanced Biology, 2024/11/01. 8(11).

8. Si, L., et al., A human-airway-on-a-chip for the rapid identification of candidate antiviral therapeutics and prophylactics. Nature Biomedical Engineering 2021 5:8, 2021-05-03. **5**(8).

9. Travaglini, K.J., et al., A molecular cell atlas of the human lung from single-cell RNA sequencing. Nature 2020 587:7835, 2020-11-18. **587**(7835).

10. Edington, C.D., et al., Interconnected Microphysiological Systems for Quantitative Biology and Pharmacology Studies. Scientific Reports 2018 8:1, 2018-03-14. **8**(1).

11. Loewa, A., J.J. Feng, and S. Hedtrich, Human disease models in drug development. Nat Rev Bioeng, 2023: p. 1–15.

12. Altulea, D., et al., What makes (hydroxy)chloroquine ineffective against COVID-19: insights from cell biology. Journal of Molecular Cell Biology, 2021 Mar 9. 13(3).

13. Yan, V.C. and F.L. Muller, Why Remdesivir Failed: Preclinical Assumptions Overestimate the Clinical Efficacy of Remdesivir for COVID-19 and Ebola. Antimicrobial Agents and Chemotherapy, 2021 Sep 17. 65(10).

14. Bein, A., et al., Frontiers | Enteric Coronavirus Infection and Treatment Modeled With an Immunocompetent Human Intestine-On-A-Chip. Frontiers in Pharmacology, 2021/10/25. 12.

15. Nawroth, J.C., et al., Breathing on chip: Dynamic flow and stretch accelerate mucociliary maturation of airway epithelium in vitro. Mater Today Bio, 2023. 21: p. 100713.

16. Hogan, Brigid L.M., et al., Repair and Regeneration of the Respiratory System: Complexity, Plasticity, and Mechanisms of Lung Stem Cell Function. Cell Stem Cell, 2014/08/07. 15(2).

17. El-Bassouny, D.R., et al., Role of nuclear factor-kappa B in bleomycin induced pulmonary fibrosis and the probable alleviating role of ginsenoside: histological, immunohistochemical, and biochemical study. Anatomy & Cell Biology. Vol. 54. 2021/12/31.

18. Okuda, K., et al., Localization of Secretory Mucins MUC5AC and MUC5B in Normal/Healthy Human Airways. American Journal of Respiratory and Critical Care Medicine, 2019 Mar 15. 199(6).

19. Sidhaye, V.K., et al., Shear stress regulates aquaporin-5 and airway epithelial barrier function. Proc Natl Acad Sci U S A, 2008. 105(9): p. 3345–50.

20. Tarran, R., et al., Normal and cystic fibrosis airway surface liquid homeostasis. The effects of phasic shear stress and viral infections. J Biol Chem, 2005. 280(42): p. 35751–9.

21. Lazarowski, E.R., et al., Molecular mechanisms of purine and pyrimidine nucleotide release. Adv Pharmacol, 2011. 61: p. 221–61.

22. Zhu, Y., et al., Baseline Goblet Cell Mucin Secretion in the Airways Exceeds Stimulated Secretion over Extended Time Periods, and Is Sensitive to Shear Stress and Intracellular Mucin Stores. PLoS One, 2015. 10(5): p. e0127267.

23. Vimalajeewa, D., et al., Virus particle propagation and infectivity along the respiratory tract and a case study for SARS-CoV-2. Scientific Reports, 2022 May 10. 12(1).

24. Sharif-Askari, N.S., et al., Airways Expression of SARS-CoV-2 Receptor, ACE2, and TMPRSS2 Is Lower in Children Than Adults and Increases with Smoking and COPD. Molecular Therapy - Methods & Clinical Development, 2020/09/11. 18.

25. Yumoto, H., et al., Sensitization of Human Aortic Endothelial Cells to Lipopolysaccharide via Regulation of Toll-Like Receptor 4 by Bacterial Fimbria- Dependent Invasion. Infection and Immunity, 2005 Dec. 73(12).

26. Monick, M.M., et al., Respiratory Syncytial Virus Up-regulates TLR4 and Sensitizes Airway Epithelial Cells to Endotoxin. Journal of Biological Chemistry, 2003/12/26. 278(52).

27. R, L., et al., Differential regulation of the transcriptomic and secretomic landscape of sensor and effector functions of human airway epithelial cells - PubMed. Mucosal immunology, 2018 May. 11(3).

28. Willett, B.J., et al., SARS-CoV-2 Omicron is an immune escape variant with an altered cell entry pathway. Nature Microbiology 2022 7:8, 2022-07-07. **7**(8).

29. Guo, Z.-y., et al., COVID-19: from immune response to clinical intervention. Precision Clinical Medicine, 2024/07/24. 7(3).

30. Z, Z., et al., Heat shock protein 90 facilitates SARS-CoV-2 structural protein-mediated virion assembly and promotes virus-induced pyroptosis - PubMed. The Journal of biological chemistry, 2023 May. 299(5).

31. Shen, Z., et al., Elucidating host cell response pathways and repurposing therapeutics for SARS-CoV-2 and other coronaviruses. Scientific Reports 2022 12:1, 2022-11-05. **12**(1).

32. R, B., et al., Persistence of viral RNA, pneumocyte syncytia and thrombosis are hallmarks of advanced COVID-19 pathology - PubMed. EBioMedicine, 2020 Nov. 61.

33. Chaudhary, S., et al., Ultrastructural study confirms the formation of single and heterotypic syncytial cells in bronchoalveolar fluids of COVID-19 patients. Virology Journal 2023 20:1, 2023-05-19. **20**(1).

34. Purwono, P.B., et al., Infection kinetics, syncytia formation, and inflammatory biomarkers as predictive indicators for the pathogenicity of SARS-CoV-2 Variants of Concern in Calu-3 cells. PLOS ONE, 2024 Apr 3. 19(4).

35. Gedda, M.R., et al., Longitudinal transcriptional analysis of peripheral blood leukocytes in COVID-19 convalescent donors. Journal of Translational Medicine, 2022 Dec 12. 20(1).

36. Kulasinghe, A., et al., Profiling of lung SARS-CoV-2 and influenza virus infection dissects virus-specific host responses and gene signatures. European Respiratory Journal, 2022- 06-02. 59(6).

37. Nyberg, T., et al., Comparative analysis of the risks of hospitalisation and death associated with SARS-CoV-2 omicron (B.1.1.529) and delta (B.1.617.2) variants in England: a cohort study. The Lancet, 2022/04/02. 399(10332).

38. Goethem, N.V., et al., Clinical Severity of SARS-CoV-2 Omicron Variant Compared with Delta among Hospitalized COVID-19 Patients in Belgium during Autumn and Winter Season 2021–2022. Viruses 2022, Vol. 14, Page 1297, 2022-06-14. **14**(6).

39. Taylor, K., et al., Clinical characteristics and outcomes of SARS-Cov-2 B.1.1.529 infections in hospitalized patients and multi-surge comparison in Louisiana. PLOS ONE, 21 Oct 2022. 17(10).

40. Rajaiah, R., et al., Differential immunometabolic responses to Delta and Omicron SARS- CoV-2 variants in golden syrian hamsters. iScience, 2024/08/16. 27(8).

41. Mohandas, S., et al., Comparative pathogenicity of BA.2.12, BA.5.2 and XBB.1 with the Delta variant in Syrian hamsters. Frontiers in Microbiology, 2023 Jun 22. 14.

